# The impact of within-host coinfection interactions on between-host parasite transmission dynamics varies with spatial scale

**DOI:** 10.1101/2023.06.21.545944

**Authors:** Shaun P. Keegan, Amy B. Pedersen, Andy Fenton

## Abstract

Within-host interactions among coinfecting parasites can have major consequences for individual infection risk and disease severity. However, the impact of these within-host interactions on between-host parasite transmission, and the spatial scales over which they occur, remain unknown. We developed and apply a novel spatially-explicit analysis to parasite infection data from a wild wood mouse (*Apodemus sylvaticus*) population. We previously demonstrated a strong negative interaction within individual hosts gastrointestinal parasites, the nematode *Heligmosomoides polygyrus* and the coccidia *Eimeria hungaryensis*, using drug-treatment experiments. Here, we find that this negative within-host interaction can significantly alter the between-host transmission dynamics of *E. hungaryensis*, but only within spatially-restricted neighbourhoods around each host. However, for the closely-related species *E. apionodes*, which experiments show does not interact strongly with *H. polygyrus*, we did not find any effect on transmission over any spatial scale. Our results demonstrate that the effects of within-host coinfection interactions can ripple out beyond each host to alter the transmission dynamics of the parasites, but only over local scales that likely reflect the spatial dimension of transmission. Hence there may be knock-on consequences of drug treatments impacting the transmission of non-target parasites, altering infection risks even for non-treated individuals in the wider neighbourhood.

## Introduction

Most individuals, among wildlife, livestock and humans in many parts of the world, are infected by multiple parasite species throughout their lives (1). It is well known that such coinfecting parasites can interact strongly with each other within individual coinfected hosts (2–7), with potentially important implications for host susceptibility, clinical disease progression, and treatment and vaccine efficacy (8). However, most attention to date has focused primarily on the mechanisms driving within-host interactions, and the effects on the individual hosts they occur within, with little understanding of the consequences these interactions have for parasite transmission between hosts, or for the role they play in driving spatiotemporal and epidemiological dynamics of infection across larger spatial scales.

Intuitively we may expect within-host coinfection interactions to impact parasite population dynamics through a process of ‘transmission modification’, whereby coinfection alters the production of parasite transmission stages, thereby modifying the risk of other hosts in the population becoming infected. Existing theory broadly supports this hypothesis, but also suggests that the nature of the scaling relationship from within-host interaction to population-level impact may be highly non-linear, dependent, at least in part, on the mechanism underlying the within-host interaction (9–11). Some evidence from natural systems supports these predictions. For example, contrasting individual- and population-level effects of helminth – bovine tuberculosis (bTB) coinfection have been predicted in African Buffalo (12), whereby anthelmintic-treatment benefited individuals through reduced bTB-induced host mortality, but this was predicted to increase bTB transmission at the population level (12). Furthermore, analyses of specific gastrointestinal nematode and protozoan infections in UK wood mice (*Apodemus sylvaticus*) find no signal of interspecific associations at the host population scale (13), even though drug treatment experiments in wild populations and controlled coinfection experiments in the laboratory clearly show the nematodes strongly suppress the protozoans within individual coinfected hosts (4,14,15). Together these results indicate that within-host coinfection interactions can have potentially counterintuitive effects on parasite transmission through the host population.

We suggest that a key aspect missing from coinfection studies to date, that prevents clear assessment of the impact of within-host interactions on between-host transmission dynamics, is spatial scale. Most studies either look exclusively at the within-host scale (e.g., through controlled infection/coinfection experiments on individual hosts), or the whole-population scale (e.g., looking for patterns of parasite co-association across the entire host population). Yet parasite transmission is an inherently spatial process: the movement of a parasite from one individual to another typically requires a degree of spatial proximity, the magnitude of which may vary with transmission mode. Hence, we may expect the effects of transmission modification due to coinfection interactions to show strong spatial structuring. To date, however, there has been no explicit empirical assessment of how within-host parasite interactions affect parasite transmission in a natural system, nor quantification of the spatial scale over which such effects occur.

We present a spatially-explicit analysis of the between-host transmission consequences of within-host interactions among three coinfecting parasites in a wild wood mouse population. Through anthelmintic drug treatment experiments in these natural populations, we have previously shown that the dominant gastrointestinal (GI) nematode, *Heligmosomoides polygyrus* (prevalence ∼50-90%), suppresses coinfections by the coccidial parasite *Eimeria hungaryensis* (Knowles et al 2013). Animals treated with the anti-nematode drug Ivermectin showed a 15-fold increase in *E. hungaryensis* oocyst shedding 1-3 weeks post-treatment, compared with untreated animals. Hence, *H. polygyrus* interacts negatively with *E. hungaryensis*, suppressing output of its infective stages from coinfected animals.

Notably, however, there was little impact of treatment on the closely related species, *E. apionodes* (Knowles et al 2013), suggesting that *H. polygyrus* has little impact on this species in coinfected mice. The hypothesised reason for this difference between the two *Eimeria* species arises from the fact the species differ in their preferential infection niches in the gut: *E. hungaryensis* and *H. polygyrus* both infect in the same part of the small intestine, specifically the duodenum, whereas *E. apionodes* infects further down the small intestine, in the lower ileum (4,15). Hence the impact of *H. polygyrus* on coinfecting *Eimeria* is highly localised within the gut, due either to competition for shared resources or physical interference (e.g., disruption of epithelial tissue integrity and/or reduction in the availability or longevity of gut epithelial cells), or localised immune responses (e.g., reduction in specific and total antibodies during coinfection (4,16)), that have little impact on *E. apionodes* further down the gut. The contrasting *H. polygyrus*-mediated coinfection effects between these two closely-related parasite *Eimeria* species with otherwise very similar life-cycles provides an ideal opportunity to compare the population-dynamic consequences of the within-host interaction. Specifically, we hypothesise that increasing prevalence of *H. polygyrus*-infected animals will, through coinfection-mediated transmission modification, reduce the local force of infection of *E. hungaryensis* sufficiently to reduce *E. hungaryensis* infection in focal animals within the neighbourhood; however, we predict to find no such effect for *E. apionodes*. We also hypothesise that any effect of transmission modification will vary with neighbourhood size around focal animals, and will be most apparent at local spatial scales where transmission of these faecal-oral gastrointestinal parasites will likely occur. We test these hypotheses through a spatially-explicit analyses of wood mouse trapping data over a range of neighbourhood sizes, and show that within-host coinfection interactions do indeed influence between-host transmission of *E. hungaryensis*, but not *E. apionodes*, but the effects for *E. hungaryensis* are strongest over relatively localised spatial scales around individual hosts.

## Materials and Methods

### Wood mouse and parasite data collection

Individual wood mice (*Apodemus sylvaticus*) were trapped every 2 weeks, using baited Sherman traps (Alana Ecology, UK; dimensions 8.9cm x 7.6cm x 22.9cm), from May to December 2012 in Haddon Wood, England (Wirral Peninsula, 51.0508ºN, 3.4980ºW). Traps were laid on a semi-permanent 110 × 110 m grid, with 2 traps every 10 m, which were checked every day for 3 consecutive days, every 2 weeks. At first capture, all mice were permanently tagged with a subcutaneous microchip transponder for identification (AVID Friend Chip). For each mouse at every capture, we took the following metrics: body length (nose tip to base of tail in mm), weight (g), sex, reproductive status and an estimate of age (see (4) for additional methodological details). At every capture, we also collected faecal samples from previously sterilised, single occupancy traps for faecal floatation and microscopic analysis to identify and quantify both helminth eggs and coccidial oocysts (measured as eggs or oocysts/gram; see (4) for details of identification and quantification of infection by these gastrointestinal parasites). As part of an on-going experiment, a subset of animals received either weight-adjusted dose of Ivermectin (an anthelmintic known to reduce nematodes: Eqvalan, 9.4 mg/kg; n=68 animals treated) or Vecoxan (an anti-coccidial treatment; n=63 animals treated) or both (n=57 animals); all other animals (n=70) received water as a control. There were 986 captures in total. All mice were released at the point of capture after handling.

### Neighbourhood Analysis: data structure

We considered each individual at a single capture point as the ‘focal’ individual in turn. Around each focal animal at each capture, we defined its neighbourhood of size *r* as all trap locations lying within *r* metres of the focal’s capture location (Fig. 1). We then determined the number of other individuals caught at traps within that neighbourhood within a specified time window prior to the capture date of the focal individual; animals caught after the focal’s capture date were ignored, as they could not have contributed to the focal individual’s infection status at that time point, and animals caught prior to the specified time window were deemed unlikely to have contributed current infection levels in the focal host. We explored sensitivity of our findings to the duration of this window by running our analyses to a range of time window sizes (17 days, 34 days, 51 days, or all previous time points prior to the focal’s capture date; date ranges are dictated by trapping frequency/experimental design). For each specified time window and neighbourhood size around each focal individual, we calculated the proportion of neighbouring hosts within that neighbourhood that were infected with the nematode *H. polygyrus*. We compiled this dataset using all captured mice as focal animals for all possible neighbourhood sizes, from *r* = 10 m (comprising the 4 closest neighbours only around each focal capture) up to *r* = 50m (approximately half the trapping grid). For some individuals and neighbourhood sizes, it was not possible to account for their entire neighbourhood as some of it lay outside the experimental grid. In these instances, we assumed that the grid was representative of the wider woodland, and as such the observed prevalence of infection within the neighbourhood on the grid reflected that of the focal’s entire neighbourhood.

**Figure 1.**
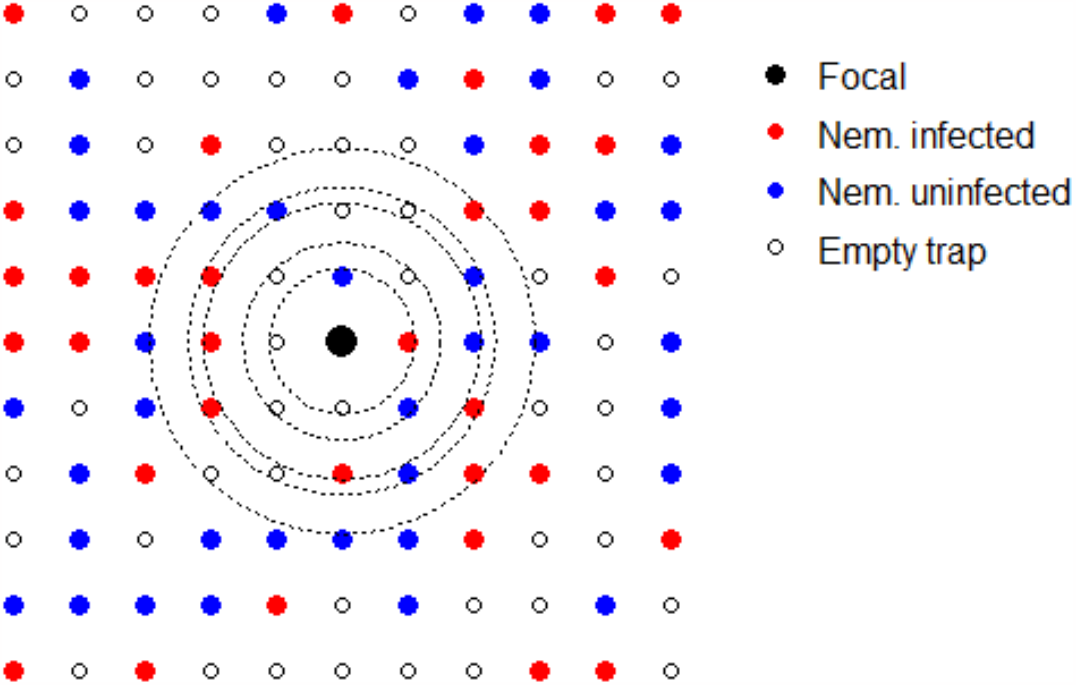
Example trapping grid with wood mouse neighbourhoods. This trapping grid shows the relative locations of traps, with the current focal animal shown in black. Neighbouring traps are colour-coded either red for an animal infected with the nematode *H. polygyrus*, blue for an uninfected animal, or blank for no capture within the specified time window of the focal’s capture. Five example neighbourhoods of increasing size around the focal are shown in dotted outline. The prevalence of nematode infection within each neighbourhood size of the focal is then calculated as the proportion of infected animals (red points) out of the total number of animals caught (red + blue points) within that neighbourhood, for the specified time window.

### Neighbourhood Analysis: statistical analyses

We analysed the above datasets for each specified time window in turn, to determine whether there was a signal of increasing neighbourhood prevalence of the nematode *H. polygyrus* on each focal individual’s infection with either *E. hungaryensis* or *E. apionodes*; a significant effect of neighbourhood *H. polygyrus* prevalence on focal *Eimeria* infection levels would suggest that the within-host effects of *H. polygyrus* coinfection modify oocyst shedding of that *Eimeria* species sufficiently to alter its within-neighbourhood force of infection. For each specified time window, and for all possible neighbourhood sizes up to 50m radius around each focal individual (*r* = 10m, 14.1m, 20m, 22.4m, 28.3m, 30m, 31.6m, 36.1m, 40m, 50m; neighbourhood sizes are dictated by trapping array), we ran Generalized Linear Mixed Effects Models (GLMMs) implemented in a Bayesian framework using stan (R packages: rstan (version 2.17.3), brms (version 2.3.0)). Each individual’s *Eimeria* infection status (either *E. hungaryensis* or *E. apionodes* in turn) was quantified as infection intensity (number of oocysts per gram of faeces among infected animals), as the response variable in a Gaussian GLMM. Our main fixed effect of interest was the neighbourhood prevalence of *H. polygyrus* (proportion of hosts infected within the specified neighbourhood size around the focal individual, within the specified time window). In all models we also controlled for the following potential confounders of the focal individual: its *H. polygyrus* infection status (positive/negative), sex (male or female) and age (adult or not). We also controlled for the total number of animals in the defined neighbourhood, and the date of capture as a 2^nd^-order polynomial to account for non-linear seasonal effects, and focal ID (unique PIT tag number) as a random effect to control for potential pseudo-replication arising from multiple captures of the same individual. We did not carry out any model reduction or simplification, to ensure consistency in model structure across all analyses, and we report the effect sizes (median GLMM coefficients, ± 95% credible intervals) for neighbour *H. polygyrus* prevalence on focal host *E. hungaryensis* or *E. apionodes* infection intensity, for each neighbourhood size and time window, while controlling for the same set of potential confounders in all analyses. Negative effect sizes would suggest that increasing prevalence of *H. polygyrus* was associated with a reduction in the intensity of *Eimeria* infection of focal individuals within the specified time window and neighbourhood size.

We note that our focus is on how *H. polygyrus* prevalence within a specified neighbourhood size relates to *Eimeria* infection levels in each focal host. As such, increasing neighbourhood sizes around each focal individual incorporated the same hosts that were found within smaller neighbourhoods around that focal individual. Hence successive neighbourhoods are not independent of each other. Due to that non-independence, we did not seek to include all possible neighbourhoods in a single analysis, with a model term describing how the effect of *H. polygyrus* changes with neighbourhood size. Rather, we conducted a series of analyses to understand how changing spatial scale altered the association between local *H. polygyrus* prevalence and focal *Eimeria* infection intensity. This approach requires multiple testing through repeated GLMM analyses. For this reason, in part, we did not evaluate statistical significance of effects through the calculation of p-values for each test. Instead we adopted the above Bayesian approach to generate a series of estimated effect sizes, which we use to infer the consequence of increasing *H. polygyrus* prevalence on focal *Eimeria* infection levels at each neighbourhood size.

To ensure that the above method is not biased to generating associations when there are none, we carried out a randomisation test, whereby we reran the analyses on a dataset that randomly assigned observed pairs of *H. polygyrus* and *E. hungaryensis* infection data from each animal to the spatial and temporal capture records to other animals in the dataset. That is, we retained individual-level *H. polygyrus* and *E. hungaryensis* associations, but detached those from their neighbourhood context.

Hence this approach tests, for realistic host spatiotemporal captures, and parasite (co)infection data, whether our neighbourhood analysis method generates spurious associations between *H, polygyrus* neighbourhood prevalence and *Eimeria* infections, even in the absence of a true relationship in the data.

### Accounting for treated animals

As mentioned above, a subset of animals in the population received the anti-parasite drugs Ivermectin (which targets nematodes) or Vecoxan (which targets coccidial parasites like *Eimeria*). We have previously shown that Ivermectin is effective in reducing the intensity of nematode infection in wild wood mice, but the effect is transient, with treated individuals being reinfected within days of treatment (4,17). For our analyses we excluded animals that had been treated with Ivermectin as focal individuals, as it may confound our results. However, they were allowed to be considered as neighbours (i.e., they do occur on the grid, and so could potentially contribute to infection of focal individuals). We have not detected any observable effect of Vecoxan on either *Eimeria* species in these wood mouse populations (18) and so, given this lack of observable effect of treatment, we included Vecoxan-treated animals as both focal and neighbouring hosts in our analyses.

All analyses were undertaken in R (version 3.5.0).

## Results

Our dataset comprised 986 captures of 252 wild wood mice with information on their *H. polygyrus* and *Eimeria* spp. infection statuses, and spatially-explicit and time-referenced information on their capture locations. Our neighbourhood analysis revealed a general reduction in individual-level infection intensity of *E. hungaryensis* with increasing *H. polygyrus* infection prevalence among all other individuals in the surrounding neighbourhood (Fig. 2). Hence, the greater the proportion of *H. polygyrus*-infected animals in a neighbourhood, the lower the average intensity of *E. hungaryensis* infections within each focal individual. Note that the statistical models controlled for the *H. polygyrus* infection status of the focals (as well as other potentially confounding factors: age, sex and time of year), and so these neighbourhood-level effects are over-and-above any direct effect of *H. polygyrus* on *E. hungaryensis* infections within the focals themselves. Notably, however, this reduction in focal *E. hungaryensis* intensity with increasing neighbourhood *H. polygyrus* prevalence was relatively localised, being most apparent at spatial scales that approximate the mean home range size of wood mice (∼15-30m radius) (19). At larger spatial scales (∼30-50m), we found no association between neighbourhood *H. polygyrus* prevalence and *E. hungaryensis* intensity in focal hosts at most time-windows (with the exception of 51 days). For all time windows, there were no detectable associations between *H. polygyrus* neighbourhood prevalence and focal *E. hungaryensis* intensity for any neighbourhood sizes >31m, consistent with our previous population-level analyses that found no associations between *H. polygyrus – E. hungaryensis* infections across the wider mouse population (13). Broadly, the neighbourhood effects were robust across all temporal scales examined, however larger spatial effects are seen at the 51-day time window. Furthermore, effect size estimates were least certain (largest 95% credible intervals) when all time points were included (Fig 1, bottom panel), suggesting that including the effects of neighbours over all time points prior to the focal’s capture may obscure the signal of the interaction effect with *H. polygyrus*.

**Figure 2.**
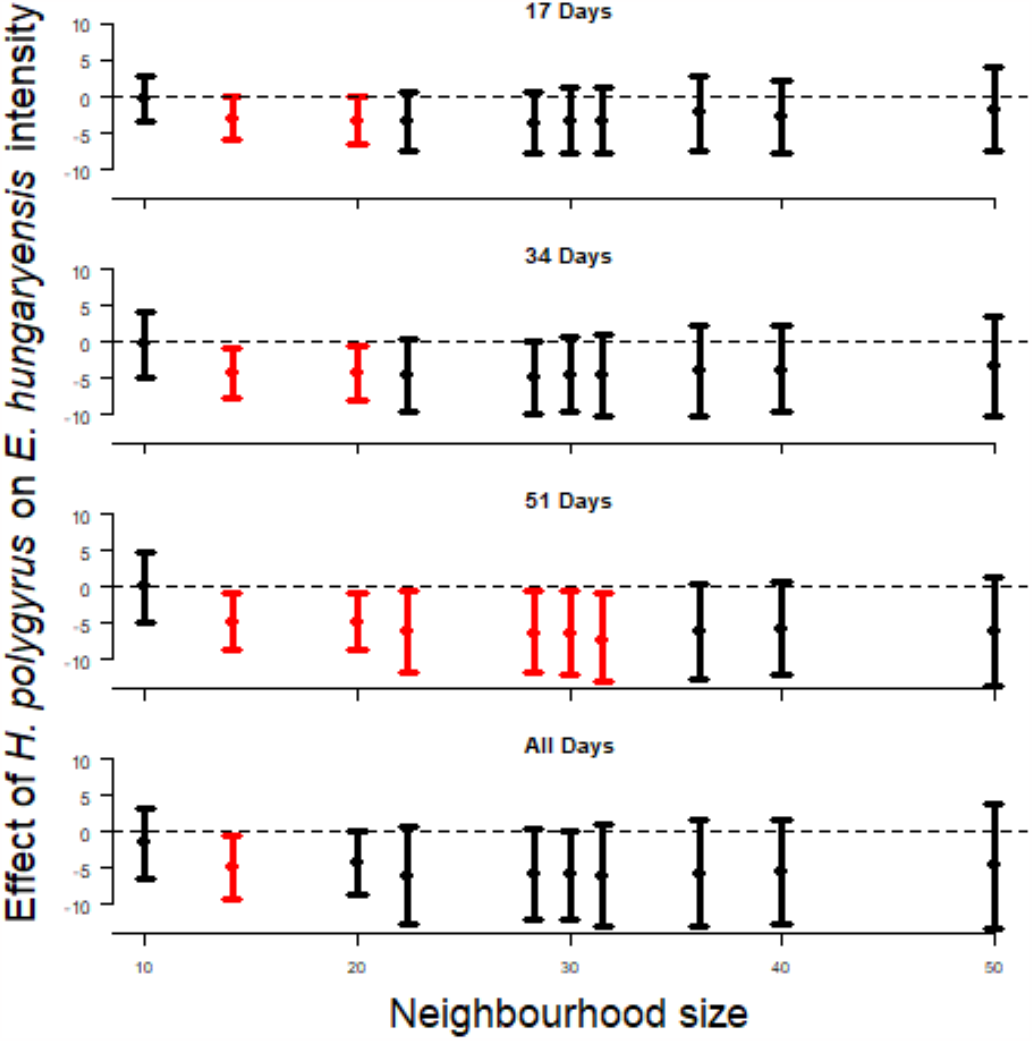
Neighbourhood analysis of parasite data from wood mice (*Apodemus sylvaticus*) showing the associations between the effect size (from GLMMs) of the neighbourhood-level prevalence of the GI nematode *H. polygyrus* on focal individual *E. hungaryensis* infection intensity (known from previous experimental perturbations (4), to undergo strong within-host coinfection interactions with the *H. polygyrus*), for increasing neighbourhood sizes (x-axes). Each panel shows results for different time windows between captures of animals in the neighbourhood and subsequent capture of the focal individual. Figures show median model estimates (points) and 95% credible intervals (envelopes) from Bayesian GLMMs of the effect of neighbourhood prevalence of *H. polygyrus* on the relevant individual-level infection metrics (intensity or probability of infection) for each of the focal parasites, at each neighbourhood size, for each time window. Red points denote neighbourhood sizes for which the 95% credible intervals do not cross 0.

The above analyses suggest that the known, negative within-host effect of *H. polygyrus* on *E. hungaryensis* oocyst shedding leaves a signal of that interaction on the transmission dynamics of *E. hungaryensis* within the local neighbourhood. Comparison with the closely-related species *E. apionodes*, which we know from previous drug-treatment experiments does not interact strongly with *H. polygyrus*, helps add veracity to these findings. As hypothesised, we see no signal of an effect of *H. polygyrus* neighbourhood prevalence on *E. apionodes* intensity at any neighbourhood size, or any time window duration (Fig. 3). Furthermore, analysis of a permuted dataset in which real mouse capture locations and timings were randomly assigned *H. polygyrus* and *E. hungaryensis* infection data, found no signal of a relationship between *H. polygyrus* neighbourhood prevalence and *E. hungaryensis* infection intensities at any spatial or temporal scales (Fig S1). Together, these neutral findings for the ‘natural control’ species *E. apionodes* and from the randomised dataset, help reassure that the findings of neighbourhood-wide suppression of *E. hungaryensis* infections were not due to a statistical artefact of the neighbourhood analysis method.

**Figure 3.**
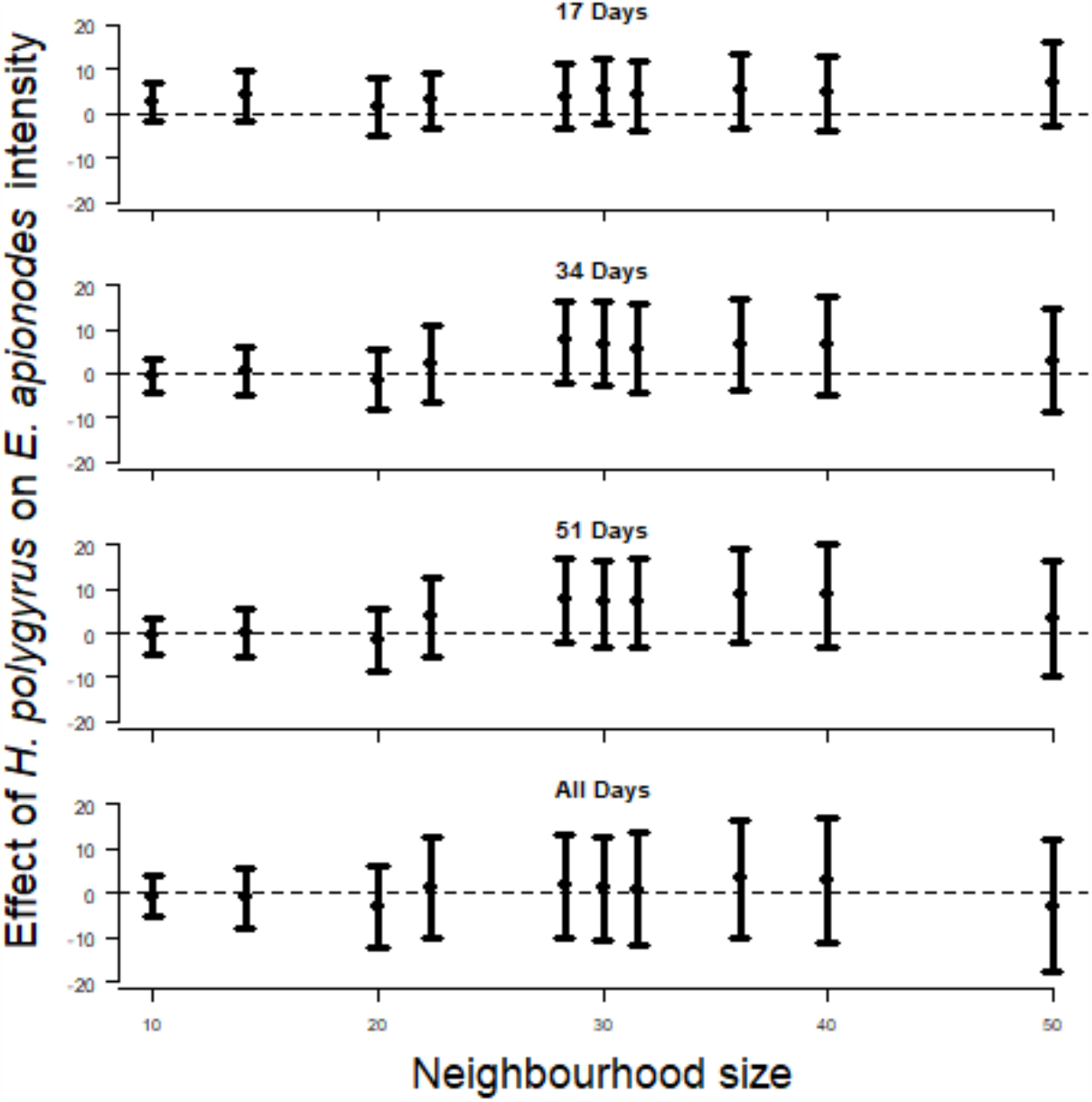
As in Fig 2, but showing results for *E. apionodes*, the species known from previous experimental perturbations (4), to not undergo strong within-host coinfection interactions with the nematode *H. polygyrus*.

## Discussion

Our study presents a spatially explicit analysis to assess how within-host interactions between coinfecting parasites alter between-host transmission dynamics across a range of spatial scales. We show that the previously-recognised negative interaction between the GI nematode *H. polygyrus* and the protozoan *E. hungaryensis* (4,15) ripples out beyond individual hosts, reducing *E. hungaryensis* transmission to neighbouring hosts. However the detectability of this effect was generally limited to neighbourhoods of ∼15-20m (with exceptions of wider detectability at certain time-windows) around focal host individuals, approximating the radius of the home range of these mice (19). Hence, the suppression of *E. hungaryensis* oocyst shedding by coinfecting *H. polygyrus* impacts the force of infection of the protozoan in the population, but only over spatial scales that represent the likely space use of each mouse. Confidence in our results is provided by the natural control of the closely-related species *E. apionodes*, which has the same transmission mode and life-cycle as *E. hungaryensis* but, due to its infection location in the mouse’s gut (found primarily lower in the small intestine), does not interact strongly with *H. polygyrus* (4). As hypothesised, our analysis found no signal of a suppressive effect of *E. apionodes* transmission by *H. polygyrus* infection over any neighbourhood size. This, together with our randomisation analysis, helps reassure that the suppressive effects seen for *E. hungaryensis* are unlikely to be artefacts of our analytical technique, and so adds further weight to the validity of the spatially-structured suppressive effects seen between *H. polygyrus* and *E. hungaryensis*. To our knowledge this is the first empirical demonstration of the occurrence and spatial extent of knock-on consequences of within-host coinfection interactions for parasite transmission in a natural host-parasite community.

Our findings have significant implications for the use of population-level analyses to screen for potential coinfection interactions between parasites. The effects we detected were highly spatially dependent, and only occurred over restricted distances within the local neighbourhood of each host; as the neighbourhood size was increased beyond ∼30m, the magnitude of the suppressive effect diminished. This lack of association between *H. polygyrus* and *E. hungaryensis* at larger spatial scales corresponds with previous analyses that have failed to detect a signal of the *H. polygyrus* – *E. hungaryensis* interaction at the level of the whole population (13). Notably, we also found that the associations between neighbourhood *H. polygyrus* prevalence and focal *E. hungaryensis* infections were least clear when we included all neighbours at any time previous to the focal individual’s current capture point (Fig 1 bottom panel). Shorter time windows likely account more accurately for time delays between oocyst shedding, development to infectiousness in the environment and subsequent exposure to focal hosts. Longer time windows will combine a wide range of shedding-to-infection processes, obscuring meaningful temporal associations between neighbour and focal infections. Overall these findings suggest that snapshots of parasite associations at the full population level (i.e., combining data over relatively large spatial and temporal scales) will likely be unable to reliably infer within-host interactions between parasites. However, examining parasite associations at smaller spatial and temporal scales, which more appropriately reflect local host movement, space use, and parasite transmission biology, will more accurately reveal underlying interactions, while not generating spurious inferences of interactions between species that do not interact strongly (e.g., *H. polygyrus* and *E. apionodes* (4)). Our approach, which explicitly considers both spatial and temporal scales, therefore has potential to identify possible coinfection interactions that otherwise are only detectable by experimental intervention. More broadly these points highlight the importance of adopting a joint spatiotemporal approach, rather than either purely spatial or purely temporal, in elucidating key aspects of parasite transmission dynamics in natural systems (19).

The spatial heterogeneities revealed here arise from two standard epidemiological features of any host-parasite system. Firstly, hosts are not homogenously mixed, for example due to territoriality, resource distribution, mating behaviours etc. Secondly, parasites are not necessarily homogenous in a host population, for example due to variations in individual host immune state and host behaviour or environmental heterogeneities which affect external parasite stages (19), creating both hot- and cold-spots of parasite occurrence even in well mixed host populations. Coupling this natural variation in parasite distribution with a non-uniform host population suggests that spatial heterogeneities are likely to be an important part of most host-parasite communities, and hence a driver of coinfection-mediated transmission modification.

Overall, we have shown that within-host interactions between coinfecting parasites affect local transmission dynamics, leaving a signal of the within-host interaction at spatial scales that ripple out beyond the individual host through a process of transmission modification, impacting infection risk and intensity for the neighbouring hosts. However, the spatial extent of these effects tends to be relatively localised, likely reflecting the spatial scale of transmission and the movement/space use patterns of the hosts. There are many human, livestock and wildlife disease systems where there are well-documented within-host interactions among coinfecting parasites, and a growing body of research has examined the consequences of those interactions for the success and impact (beneficial or detrimental) of disease treatment approaches on individual host health (20,21). A major implication of our work is that there could be knock-on, between-host consequences of such treatments, particularly in communities experiencing high coverage mass drug administration, for localised transmission dynamics of non-target parasites, with implications for altered infection risk among non-treated individuals in the wider neighbourhood.

## Acknowledgments

We would like to thank those involved in the fieldwork, which was carried out by Becci Barber, Łukasz Łukomski, Steve Price, Godefroy Devevey and Susan Withenshaw, along with the assistance of many undergraduate helpers from the University of Liverpool. We thank the landowners for permission to carry out the work on their land. Finally, we would like to extend our thanks to the Ecology, Evolution and Genomics of Infectious Diseases group at the University of Liverpool, and the Edinburgh Glasgow and Liverpool Infectious Disease Ecology group for constructive comments and discussion throughout this project.

## Funding

The work was funded by NERC Grants NE/I024038/1 and NE/I026367/1 awarded to AF and ABP. SPK’s PhD was funded by the NERC ACCE Doctoral Training Partnership; ABP was funded by a Wellcome Trust Strategic Grant (095831) and a University of Edinburgh Chancellors Fellowship.

## Author Contributions

Conceptualisation: SPK & AF. Formal analysis: SPK & AF. Writing – original draft: SPK & AF. Writing – review & editing: SPK, AF & ABP. Funding acquisition and project administration: AF & ABP.

## Data and materials availability

https://github.com/shaunkeegan/coinfection_spatial_scaling_paper

## Notes

https://github.com/shaunkeegan/coinfection_spatial_scaling_paper

### Competing Interest Statement

The authors have declared no competing interest.

