## Supplementary Figure 1 for "The impact of within-host coinfection interactions on between-host parasite transmission dynamics varies with spatial scale"

**Fig S1**. Results from neighbourhood analysis showing the associations between the effect size (from GLMMs) of the neighbourhood-level prevalence of the GI nematode *H. polygyrus* on focal individual *E. hungaryensis* infection intensity for increasing neighbourhood sizes, applied to a dataset that randomly assigned observed pairs of *H. polygyrus* and *E. hungaryensis* infection data from each animal to the spatial and temporal capture records to other animals in the dataset. Each panel shows results for different time windows between captures of animals in the neighbourhood and subsequent capture of the focal individual. Figures show median model estimates (points) and 95% credible intervals (envelopes) from Bayesian GLMMs of the effect of neighbourhood prevalence of *H. polygyrus* on the relevant individual-level infection metrics (intensity or probability of infection) for each of the focal parasites, at each neighbourhood size, for each time window.

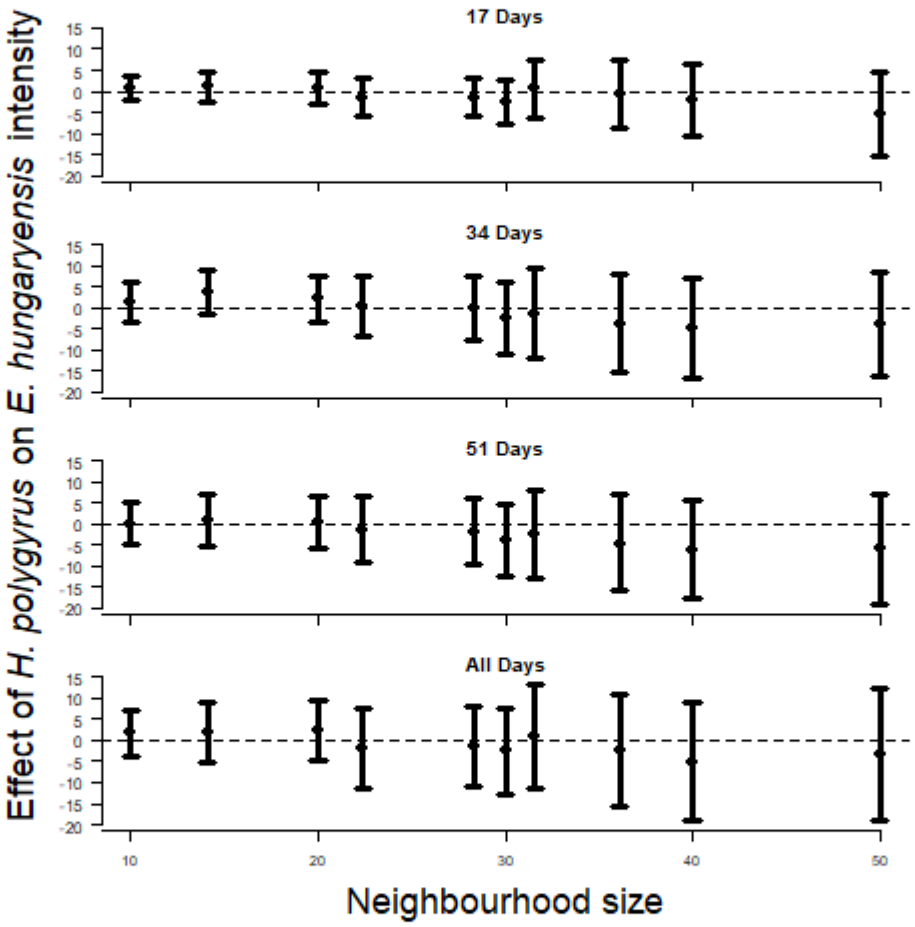
